# Expanding the Chinese hamster ovary cell long non-coding RNA transcriptome using RNASeq

**DOI:** 10.1101/863241

**Authors:** Krishna Motheramgari, Ricardo Valdés-Bango Curell, Ioanna Tzani, Clair Gallagher, Marina Castro Rivadeneyra, Lin Zhang, Niall Barron, Colin Clarke

## Abstract

Our ability to study Chinese hamster ovary (CHO) cell biology has been revolutionised over the last decade with the development of next generation sequencing and the publication of reference DNA sequences for CHO cells and the Chinese hamster. RNA sequencing has not only enabled the association of transcript expression with bioreactor conditions and desirable bioprocess phenotypes but played a key role in the characterisation of protein coding and small non-coding RNAs. The annotation of long non-coding RNAs, and therefore our understanding of their role in CHO cell biology, has been limited to date. In this manuscript, we use high resolution RNASeq data to more than double the number of annotated lncRNA transcripts for the CHOK1 genome. In addition, the utilisation of strand specific sequencing enabled the identification of more than 1,000 new lncRNAs located antisense to protein coding genes. The utility of monitoring lncRNA expression is demonstrated through an analysis of the transcriptomic response to a reduction of cell culture temperature and identification of simultaneous sense/antisense differential expression for the first time in CHO cells. To enable further studies of lncRNAs, the transcripts annotated in this study have been made available for the CHO cell biology community.

---

In recent years, successful cell line engineering strategies to enhance the production of recombinant therapeutic proteins in Chinese hamster ovary (CHO) cells through the modulation of mRNA and non-coding RNAs, such as microRNAs (miRNAs), have been reported by academic and industrial research groups (Fischer, Handrick, & Otte, 2015). As new avenues for genetic manipulation continue to be explored, long non-coding RNAs (lncRNAs), defined as transcripts longer than 200 nucleotides that do not encode a protein, have emerged as a potential route for rational design of CHO cell factories (Vito & Smales, 2018). These non-coding RNAs elicit their function through a diverse array of mechanisms including recruiting epigenetic modifying complexes to genomic loci or acting as mimics that sequester proteins or miRNAs (Fang & Fullwood, 2016). lncRNAs have been shown to interact with RNA, DNA or protein and play regulatory roles in the transcription and translation of nearby (*cis*) or distant (*trans*) genes. Functional studies in other species have revealed the importance of lncRNAs in cellular processes such as proliferation (Liu et al., 2017) and the interest in manipulating lncRNAs for improving CHO cell bioprocess performance has increased. A recent study has shown that sequence elements can also be derived from long coding RNA and be utilised to build constructs to increase recombinant protein production (Patrucco et al., 2015).

Despite the availability of several reference genomes for CHO cells (Lewis et al., 2013; Rupp et al., 2018; Xu et al., 2011) and tremendous improvements in transcriptome characterisation, lncRNAs remain poorly annotated severely limiting our ability to associate these molecules with bioprocess phenotypes, and consequently, their utility for cell line engineering. Of the three reference genomes currently available through Ensembl (release 98), the CHOK1 genome contains the largest number of annotated lncRNA genes, yet this number represents only 19% and 14% of the total number of mouse (v23) and human (v32) lncRNAs annotated in GENCODE respectively (Frankish et al., 2019). The poor conservation of lncRNAs across species and our lack of understanding of their defining sequence features in comparison to protein coding genes present considerable challenges. Next generation sequencing has, however, proven to be an important tool for lncRNA discovery in non-model organisms (Uszczynska-Ratajczak, Lagarde, Frankish, Guigó, & Johnson, 2018). In this manuscript, we describe the utilisation of high-resolution RNA sequencing (RNASeq) to significantly expand the CHO cell lncRNA landscape and subsequently demonstrate that changes in the bioreactor environment can alter the expression of these molecules.

The RNASeq data utilised for this analysis was generated as part of a study in our laboratory to advance our understanding of the transcriptomic response to decreasing CHO cell culture temperature (i.e. “temperature shift”) (see Tzani *et al*. submitted to Biotechnology & Bioengineering for details). Briefly, the experimental design of the study involved growing 8 biological replicates of a monoclonal antibody (mAb) producing CHOK1 cell lines for 48hrs at 37°C. The temperature of 4 replicates was then reduced to 31°C while the remaining 4 replicates were maintained at 37°C before all samples were submitted for transcriptomic analysis at 24hrs post temperature shift. The transition to sub-physiological cell culture temperature resulted in a reduction of growth rate, alteration of extracellular metabolite concentration as well as widespread changes in mRNA abundance and splicing (see Tzani *et al*. submitted to Biotechnology & Bioengineering).

The RNASeq data acquired is ideally suited for lncRNA discovery and expression analysis as a result of (1) the elimination of rRNA from total RNA prior to library preparation via subtractive hybridisation rather than enriching for RNAs with a polyA tail, (2) the preparation of sequencing libraries using a strand-specific method to capture lncRNAs expressed from the opposite strand to protein coding genes (i.e. antisense) and (3) capturing >50 million 150bp paired-end reads for each sample to ensure sufficient sampling depth for accurate detection and assembly of lower abundance lncRNA transcripts. To identify lncRNAs from these data a rigorous computational pipeline incorporating transcriptome assembly, lncRNA discovery methods, comparative genomics and comparison to coding and non-coding sequence databases was implemented (Figure 1).

**Figure 1:**
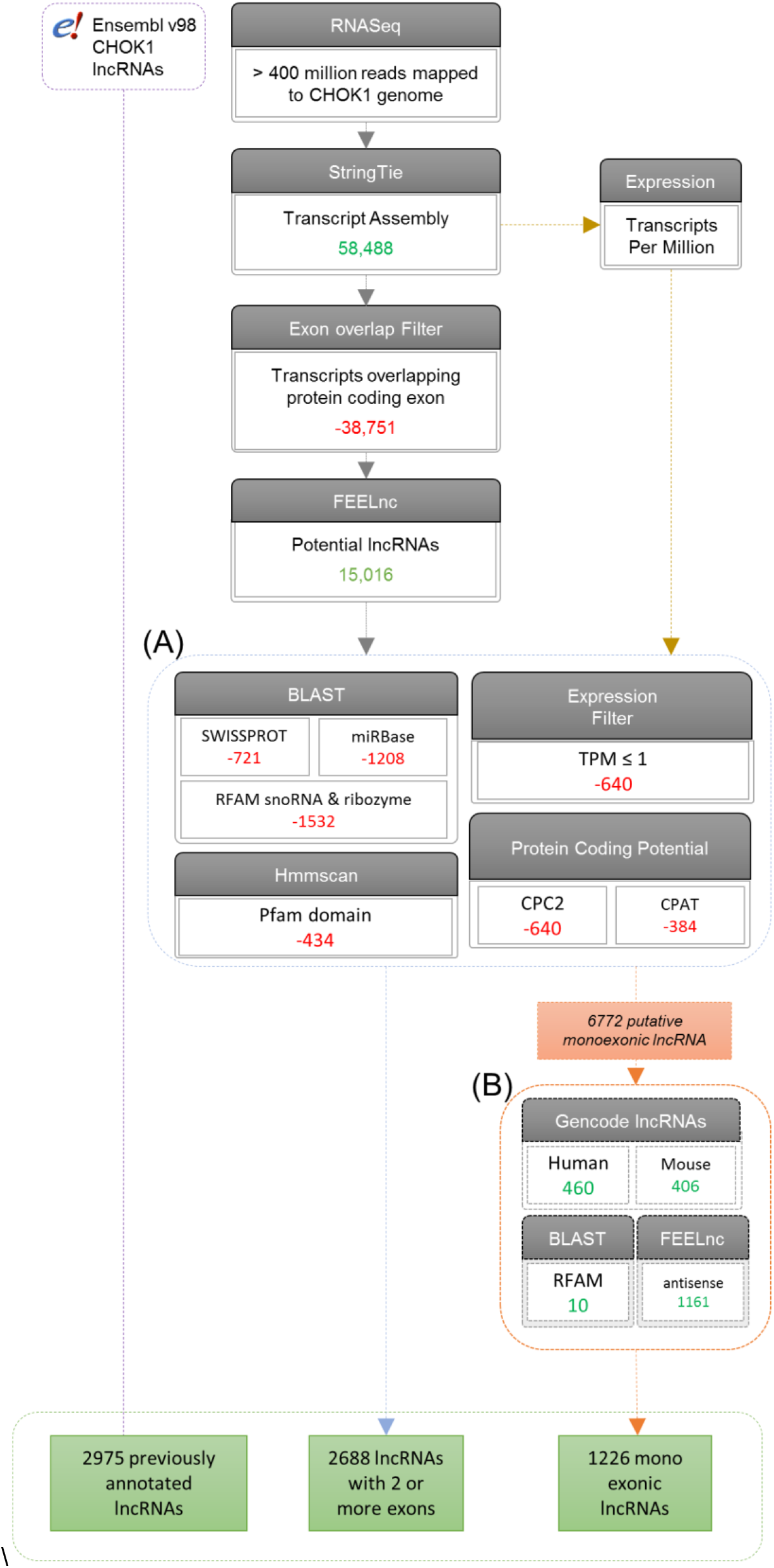
Overview of the CHO cell long non-coding RNA annotation pipeline. To accurately identify lncRNAs from the RNASeq data high quality reads mapped to the Ensembl CHOK1 genome were utilised to construct a genome guided assembly using StringTie. All transcripts that intersected with an exon of annotated protein coding genes were eliminated before FEELnc was utilised to identify putative lncRNAs. **(A)** The first filtering stage removed the FEELnc predicted lncRNA transcripts that had significant hits with non-lncRNA transcripts or protein databases. In addition, transcripts with TPM ≤ 1 or were not predicted as non-coding by CPC2 and CPAT were also removed. **(B)** The second classification stage was utilised to retain only those single lncRNAs that were orthologous with human or mouse, present in RFAM or were found on the opposite strand to an annotated protein coding gene. In total, 3,599 lncRNAs not annotated in ENSEMBL were identified bringing the total number of lncRNAs annotated for CHOK1 to 6,598.

During the initial stage of bioinformatics analysis, >400M combined high quality mappable RNASeq reads from the 8 samples were utilised to assemble >58,000 transcripts using StringTie (Pertea et al., 2015). Following the removal of >38,000 transcripts intersecting same-strand exons of protein coding genes, 19,737 transcripts were analysed using the FLExible Extraction of Long non-coding RNAs (FEELnc) lncRNA discovery algorithm (Wucher et al., 2017). FEELnc first eliminates transcripts shorter than 200 nucleotides before assigning a protein coding potential score to the remaining transcripts (the likelihood a particular transcript encodes a protein coding gene). The 15,016 putative lncRNAs identified by FEELnc were further reduced by retaining only those also determined to be non-coding by both the CPC2 and CPAT methods, found not to contain a PFAM protein domain and did not have either a significant BLAST hit against SWISSPROT, miRBase or RFAM snoRNA and ribozyme families (Figure 1A). Of the 9,410 transcripts remaining after this first-pass filtering stage, 2,688 lncRNAs with 2 or more exons were identified while the remainder were putative single exon lncRNAs. To increase the stringency of our analysis and eliminate potential false positives, only the 1,266 monoexonic lncRNAs that were antisense to protein coding gene, present in RFAM or orthologous to GENCODE annotated lncRNAs in either human or mouse were retained (Figure 1B). This analysis has resulted in the identification of 3,588 novel lncRNA transcripts and reclassification of 35 RNAs annotated as miscRNAs or pseudogenes in Ensembl (for instance, 5 of these RNAs fall below the 200bp definition of a long non-coding RNA, but were extended in our transcriptome assembly). These new lncRNAs, when combined with those currently available in Ensembl, bring the total number annotated for the CHOK1 cell line to 6,598 lncRNAs.

Comparison of the expanded lncRNA transcriptome to protein coding genes revealed that, as others have observed previously (Derrien et al., 2012), lncRNAs are generally shorter (Kolmogorov-Smirnov (KS) test p-value = < 2 × 10^−16^) (Figure 2A), less abundant (KS test p-value = < 2 × 10^−16^) (Figure 2B), and have a lower GC content than mRNAs (KS test p-value = < 2 × 10^−16^)(Figure 2C). CHO cell lncRNA genes also tended to be comprised of fewer exons (Figure 2D) and express fewer isoforms (Figure 2E) in comparison to protein coding genes, although the difference is less pronounced than observations in human and mouse due to comparatively poorer annotation of CHO cell mRNA isoforms. Intergenic lncRNAs (defined as those > 100kb from a protein coding gene) were found to be the most prevalent (n=1,982) following analysis of lncRNA genomic organisation with respect to protein coding genes (Figure 2F). In addition, a dramatic increase in the number of antisense (defined as overlapping a protein coding gene on the opposite strand of DNA) (n=1,376) and antisense divergent lncRNAs (defined as ≤ 2kb upstream of the protein gene) was identified (n=359).

**Figure 2:**
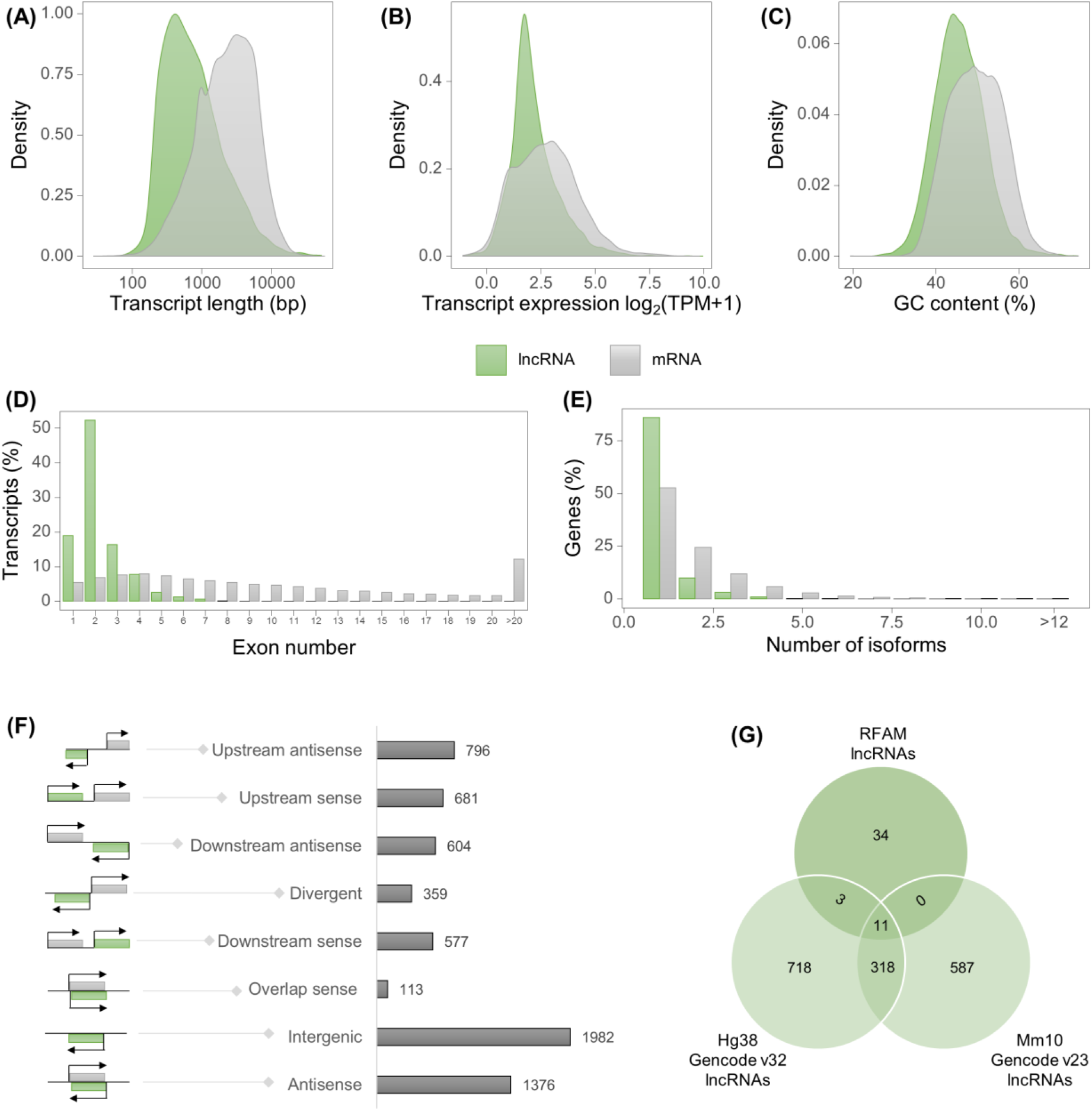
Comparison of lncRNA transcripts to annotated protein coding genes. Following comparison of the 6,598 lncRNAs annotated for CHOK1 cells to mRNA transcripts annotated in ENSEMBL, lncRNAs tend to be **(A)** shorter, **(B)** expressed at lower abundance and of generally **(C)** lower GC content. In addition, lncRNA transcripts had **(D)** a lower number of exons and **(E)** transcripts per gene. Following the determination of CHOK1 lncRNAs with respect to protein coding genes **(F)** intergenic and antisense lncRNAs were the two most prevalent classes. Sense overlap lncRNAs are likely to be underestimated due to the limitations of using RNASeq and further methods will be required to accurately annotate this type of lncRNA. The lower cross species conservation of lncRNAs is **(G)** indicated by a small number of transcripts returning significant hits from following a BLAST search against RFAM in comparison to the number of syntenic lncRNAs identified in the human and mouse genomes.

The least frequent class was found to be sense overlapping (n=113) although the conservative approach to annotation taken in this study will undoubtedly have decreased the number of these lncRNA identified. 13.8% and 15.9% of lncRNAs were found to be orthologous to lncRNA gene loci in human and mouse respectively (Figure 2G). While well-known genes including *Malat1*, had significant BLAST hits against RFAM, <1% CHOK1 lncRNAs (both novel and in Ensembl) were identified confirming the lower cross-species conservation of lncRNA sequences.

Following completion of the annotation phase, we sought to determine if long non-coding RNA abundance was changed following temperature shift of a mAb producing CHO cell line. For this analysis we utilised replicate data from both conditions and performed a gene-level count based differential expression analysis for the complete StringTie transcriptome assembly (lncRNAs and other RNAs were included in this step). The abundance of 400 lncRNAs (223 upregulated, 177 downregulated) were found to be significantly altered by ≥ ±1.5 fold change (FC) 24hrs after the reduction of cell culture temperature to 31°C (Benjamini Hochberg (BH) adjusted p-value < 0.05) (Figure 3A) (Table S1). The most frequent class of differentially expressed lncRNAs were intergenic RNAs (n=114), followed closely by antisense lncRNAs (n=94). Of the 327 differentially expressed lncRNAs unannotated in Ensembl, 176 orthologs were identified from GENCODE lncRNAs annotations for either the human or mouse genome.

**Figure 3:**
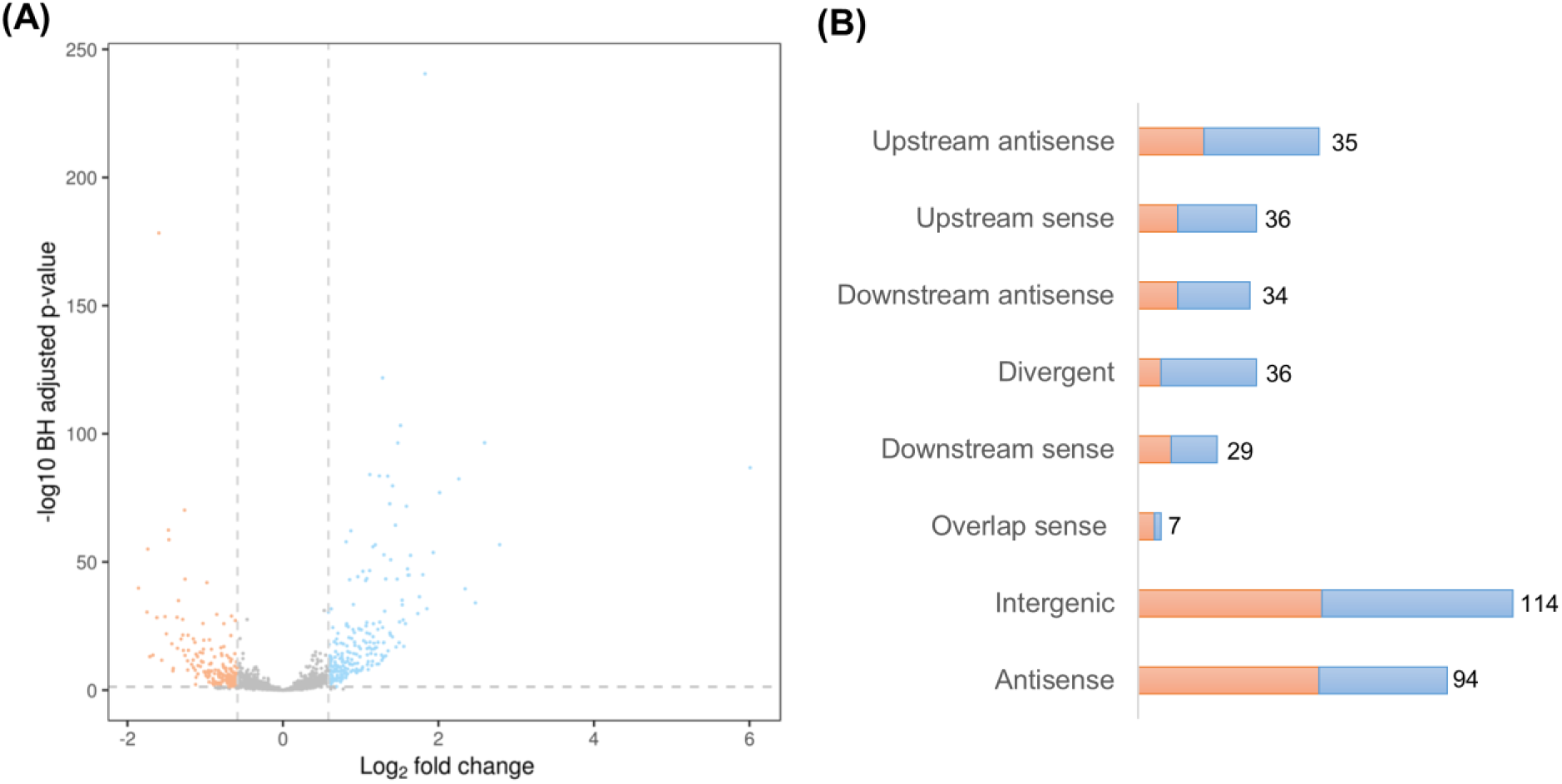
CHO cell temperature shift induces differential expression of lncRNAs. Following a reduction of cell culture temperature **(A)** the abundance of 400 lncRNAs changed were significantly altered 24hrs post-temperature shift with **(B)** 94 antisense lncRNAs found to be differentially expressed.

Next, we compared the 400 differentially expressed lncRNAs to the 1,317 differentially expressed protein coding genes identified from the same dataset using an identical count-based gene level analysis (see Tzani *et al*.). Twenty-one differentially expressed protein coding genes were found to have to have an overlapping antisense lncRNA or antisense lncRNA within 2kb upstream (divergent) that was aslo differentially expressed. 10 of these sense/antisense protein-coding/lncRNA gene pairs were found to be correlated (i.e. both the lncRNA and mRNA were upregulated) while the remaining 11 were found to be anticorrelated (i.e. the lncRNA was downregulated while the mRNA was upregulated). While these positively and negatively correlated differences in expression for protein coding and lncRNA gene pairs are certainly intriguing, these changes could occur independently (Goyal et al., 2017) and further research is required to confirm regulatory interactions.

**Figure 4:**
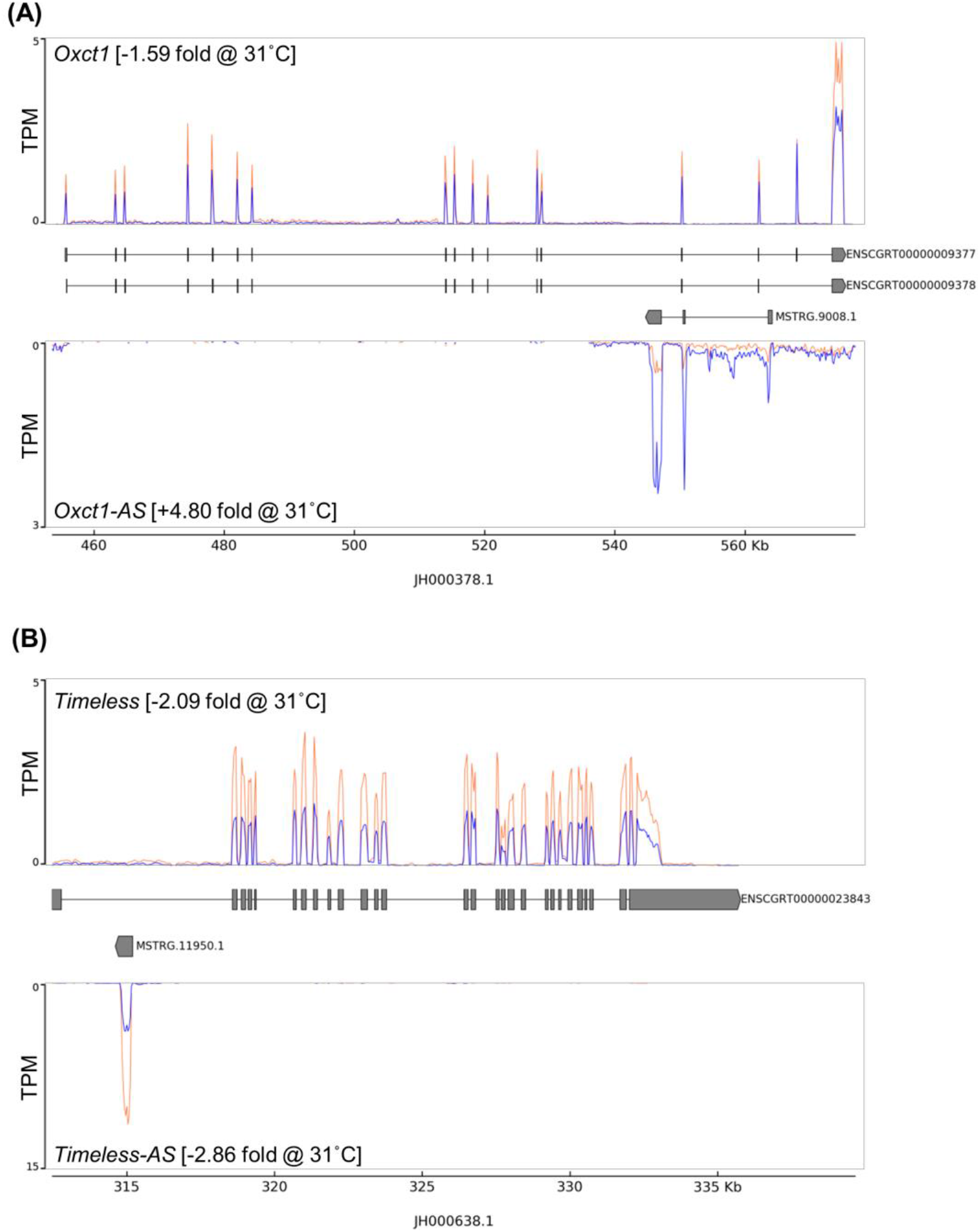
Differential expression of antisense CHO cell lncRNAs can be negatively or positively correlated to the partner mRNA found on the opposite strand. The mean TPM RNASeq coverage of the 4 biological replicates for the temp shifted (blue) and non-temperature shifted samples (orange) The **(A)** *Oxct1* gene locus is shown for both the sense and antisense strands. The expression of *Oxct1* decreases following the transition to 31°C while the expression of the *Oxct1-AS* lncRNA increases. In contrast **(B)** the *Timeless* mRNA and the *Timeless-AS* lncRNA were both found to decrease following temperature shift

In summary, the results of this study have significantly expanded the number of lncRNAs annotated for CHO cells, We have demonstrated that that lncRNA expression is altered upon a reduction of cell culture temperature and in some cases that both protein coding genes and lncRNA located in close proximity on the antisense strand can undergo correlated or anticorrelated changes in expression. More than 80% of the differentially expressed lncRNAs identified in this study are not currently annotated in Ensembl highlighting the importance of improving the characterisation of these molecules if we are to understand the role of lncRNAs in CHO cells during the production of recombinant therapeutic proteins. While much work remains to uncover the role of these molecules this study is an important step towards unlocking the potential of lncRNA for rational genetic engineering of CHO cell lines.

## 1 Materials and Methods

### 1.1 Cell culture, library preparation and RNA sequencing

The cell culture protocol, profiling of extracellular metabolites, library preparation and RNA sequencing are described in detail in Tzani *et al.* 2019 (submitted to Biotechnology and Bioengineering).

### 1.2 Data pre-processing, mapping and transcriptome assembly

Raw read data was trimmed to remove low quality bases and adaptor contamination using cutadapt (Martin, 2011) v1.18 and Trimmomatic v0.36 (Bolger, Lohse, & Usadel, 2014) and aligned to the ENSEMBL v98 CHOK1 reference genome using STAR v2.7.2d (Dobin et al., 2013). StringTie v2.0.3 (Pertea et al., 2015) (minimum junction coverage = 5) was utilised to construct a genome guided transcriptome from the aligned RNASeq data. The transcripts per million (TPM) expression value was calculated using StringTie “-e” option.

### 1.3 Long non-coding RNA annotation

#### Transcriptome assembly filtering

Those transcripts intersecting a same strand exon of an annotated protein coding gene were removed from the transcriptome assembly using BEDtools v2.25.0 (Quinlan & Hall, 2010).

#### LncRNA discovery, calculation of protein coding potential and classification

Long non-coding RNAs were initially identified following removal of transcripts < 200bp and followed by utilisation of the FEELnc software (Wucher et al., 2017). FEELnc was also used for initial classification of lncRNAs with respect to annotated protein coding genes. This classification was simplified into intergenic (>100kb from nearest protein coding genes), divergent (upstream antisense ≤ 2kb from nearest), antisense (overlapping a protein coding gene), downstream antisense, upstream antisense, sense overlapping, upstream sense, downstream sense. Additional assessment of the likelihood of each transcript encoding a protein was performed using CPC2 v0.1 (Kang et al., 2017) and CPAT v1.2.4 (Wang et al., 2013).

#### Identification of potential open reading frames in lncRNAs

TransDecoder v5.5.0 (Haas et al., 2013) was used to translate open reading frames (ORFs) longer than 100 amino acids.

#### Comparison of candidate lncRNAs to RNA and protein databases

Nucleotide and transdecoder ORF protein sequences were searched against the SWISSPROT (UniProt Consortium, 2019) (release version, July 3, 2019), miRBase (Kozomara, Birgaoanu, & Griffiths-Jones, 2019), RFAM (Kalvari et al., 2018) (lncRNA, snoRNA and Ribozyme) databases using BLAST. For all searches those hits with an expect value < 1 × 10^−5^ were considered significant. In the case of RFAM snoRNA and ribozymes searches the BLAST percentage identify value was increased to 95%.

#### Protein domain prediction

Protein domains in TransDecoder ORFs from candidate were predicted with hmmscan (Eddy, 2011) with the PfamA (El-Gebali et al., 2019). Pfam hits with an expect value < 1 × 10^−5^ were considered significant.

#### Determination of CHO cell lncRNA synteny for human and mouse

To determine if lncRNA genes were syntenic for GENCODE annotated human (v32) and mouse lncRNA (v23) lncRNA sequences were first mapped to the UCSC CHO cell genome sequence (criGriChoV1) using the GMAP algorithm (Wu & Watanabe, 2005). The UCSC genome browser (Haeussler et al., 2019) liftOver tool and prebuilt chain files linking the CHOK1 genome to human and mouse genomes were used to identify syntenic regions and subsequently GENCODE lncRNA annotations in the hg38 and mm10 genome assemblies.

### 1.4 Differential expression analysis

The number of RNASeq reads aligning to each gene in the (unfiltered) StringTie assembly was counted using HTSeq (Anders, Pyl, & Huber, 2015) using strand-specific parameters. The DESeq2 (Love, Huber, & Anders, 2014) method was used to identify differential expression between the NTS and TS sample groups [27, 28]. Those lncRNA genes with a DESeq2 base mean ≥ 100, an absolute fold change ≥ 1.5 and a Benjamini-Hochberg adjusted p value < 0.05 were considered significantly differentially expressed.

### 1.5 Reproducibility of the lncRNA annotation pipeline and RNASeq analysis

Upon acceptance of this manuscript the code used to perform lncRNA annotation and differential expression analysis will be made freely available https://github.com/clarke-lab/cho_cell_lncRNA. The RNASeq data has been deposited to NCBI SRA (PRJNA593052).

## Abbreviations

CHO: Chinese hamster ovary
lncRNA: long non-coding RNA
RNASeq: next generation RNA sequencing
BH: Benjamini Hochberg
FDR: false discovery rate
KS: Kolmogorov–Smirnov
mAb: monoclonal antibody

## Acknowledgements

The authors gratefully acknowledge funding from and Science Foundation Ireland (Grant references: 13/SIRG/2084, 15/CDA/3259 and H2020 Marie Sklodowska-Curie (Grant agreement No: 642663).

## Supplementary Tables

**Table S1: Differentially expressed lnRNA genelist.** 400 lncRNA genes were found to be differentially expressed upon comparison of the NTS and TS sample groups. The lncRNA gene ID is shown (Ensembl or StringTie assigned) along with the baseMean of DESeq2 normalised counts, log2 p-value and BH adjusted p-value for each DE lncRNA gene. Where a lncRNA was found to overlap with GENCODE annotated lncRNA the human and/or mouse Ensembl ID and gene symbol are shown. For all non-intergenic lncRNAs the “best” FEELnc classified partner CHO protein coding gene Ensembl ID, Entrez ID and description are also provided. Where the partner protein coding gene was found to be differentially expressed the fold change and adjusted p-value calculated in Tzani et al. 2019 (submitted to Biotechnology & Bioengineering).

Download link: https://app.box.com/s/4x00cj97teka5yu0w8itmsr48s6cz4jq

